# Cortical high frequency oscillations reflect encoding and retrieval of specific word concepts

**DOI:** 10.64898/2026.09.11.750906

**Authors:** Jan Cimbalnik, Sathwik Prathapagiri, Luis F. Sarmiento-Rivera, Lenka Jurkovicova, Martin Kojan, Pavel Daniel, Robert Roman, A. Czyzewski, M. Brazdil, Michal T. Kucewicz

## Abstract

High gamma and ripple frequency oscillations are engaged in encoding and recall of memory traces. Tracking specific traces with their signature neural activities across time and various tasks remains a major challenge. Using intracranial electrode recordings from epilepsy patients, we detected bursts of high-frequency oscillations in response to viewing and remembering the same common nouns in two different tasks repeated on subsequent days. We found the lowest word response selectivity of approximately 40% in the visual sensory areas and increasingly higher selectivity in the more anterior associational regions. The same word sets identified as preferred in a screening task triggered selective responses in another task during visual presentation or free recall with no sensory stimulation in the same session, and on the following day. We demonstrate that persistent reactivation of large-scale neuronal assemblies underlying particular concepts is reflected in bursts of high-frequency oscillations in the human cortex.

## Introduction

Neural activities underlying concept representations in the human mind can be directly studied using intracranial electrophysiological recordings of single neuron spiking and local field potential (LFP) oscillations [1–6]. Concept cells are a well-known example of single neuron spiking in response to various multimodal representations of the same object or place [7], as well as other cells encoding specific episodes in memory [8]. LFP oscillations are also known to be involved in encoding concept representations [9]. Gamma oscillations have been originally associated with binding coherent object representations across the visual cortex [10–12]. They were subsequently found across widespread sensory and association areas, detected as discrete bursts of oscillations spanning a broad frequency range [13–15]. More recently, coincident bursting of high gamma and ripple frequency oscillations co-occurring globally across multiple cortical areas has been proposed as a general mechanism for integrating information to bind anatomically distributed concept representations [16–19]. Hence, encoding of concepts in the brain is distributed, ranging from local spiking of individual neurons through to global cortical bursts of temporally coordinated oscillations.

High frequency oscillations (HFOs) thus offer an ideal activity for studying encoding and recall of concepts at the large-scale of electrophysiological activities. Pathological as well as physiological HFOs are detected across a broad 60-500 Hz frequency range of the LFP spectrum on both micro– and macro-scale [20–23]. These comprise bursts of oscillations at high gamma, ripple or fast ripple frequencies, which are estimated to be generated within volumes as small as a single cortical column based on micro– and macro-contact electrode recordings [21,24–26]. They effectively bridge or combine the selectivity of single neuron spiking and integrated information from coordinated bursts of neuronal assemblies [22,27]. Sequences of neuronal assembly spiking selective for particular word concepts were reported at the times of ripple HFOs during encoding and retrieval of associated word pairs [28]. Concept cells for objects and places have recently been shown to coordinate their spiking to hippocampal ripple HFOs as new object-place associations are formed [29,30]. Therefore, HFOs are thought to contain specific information about the encoded or retrieved concept representations. The degree of this selectivity compared to single neuron spiking remains underexplored [31].

What we know is that recall of particular memory items is preceded by hippocampal and cortical HFO bursts [18,25,28,32–34], which are also observed during encoding and reactivation of the remembered pictures, words or movie scenes. Neural activity present during memory encoding and reactivated before retrieval of that memory trace fulfills the criteria for an engram [35,36]. We have recently proposed HFOs as electrophysiological substrates of engrams [22]. Concepts associated with encoding and recall of specific memory traces would be reflected in the global HFO bursts, binding on their multisensory feature representations [16,18]. These global discharges engage up to half of the recorded cortical areas [18], corresponding to the proportions of cell populations engaged in encoding and recall of engrams [37]. How selective are individual HFO bursts recorded from a given cortical site to particular concepts or items remembered? And how stable are these at a given site across various tasks and time in general?

Previous studies have investigated HFOs typically within a single session of one task paradigm, limiting the ability to address their selectivity and temporal stability. Here we took advantage of unique intracranial recordings from a battery of tasks using the same lists of common nouns [38,39], which enabled us to study responses to the same words encoded and recalled in different tasks and repeated over subsequent experimental sessions. We hypothesized that at any one recording site HFO bursts will be consistently induced in response to one or more words encoded or recalled in the same task (1), in the same session but different tasks (2), and when the tasks are repeated on the subsequent day (3). Our goal was to determine how selective for particular word concepts and persistent across time are the HFO responses detected with standard intracranial macro-contact electrodes at a given cortical site.

## Results

### Word encoding and recall consistently induce physiological high-frequency oscillations

We detected 4,456,350 distinct bursts of HFOs in a large international dataset of intracranial recordings from epilepsy patients performing a battery of memory and cognitive tasks [38,39]. The detections were made in 29 patients during at least one run of Free Recall (FR) and Word Screening (WS) task (Table 1) on two consecutive days. Both tasks used the same pool of 180 common nouns, which were drawn from a standard English dataset and translated into the patients’ native languages (Czech or Slovak). These stimuli were presented for memory encoding and then freely recalled in the FR task and used for maintained attention and immediate recall during the WS task (Fig. 1a). Each subject completed one run of the FR task during the first experimental session and typically concluded the day with a run of the WS task. We repeated the session using the same word lists on the following day (Fig. 1b) to test HFO responses to specific words across different runs, assuming that the same neuronal assemblies recorded from a given electrode contact respond selectively to the presentation and recall of particular words in both tasks (Fig. 1c). Given the size of each macro-contact, we presumed that the word lists would activate multiple distinct assemblies that would generate HFOs within the same sampling field. Thus, we expected HFO responses to multiple words from the pool on any one electrode site.

**Figure 1:**
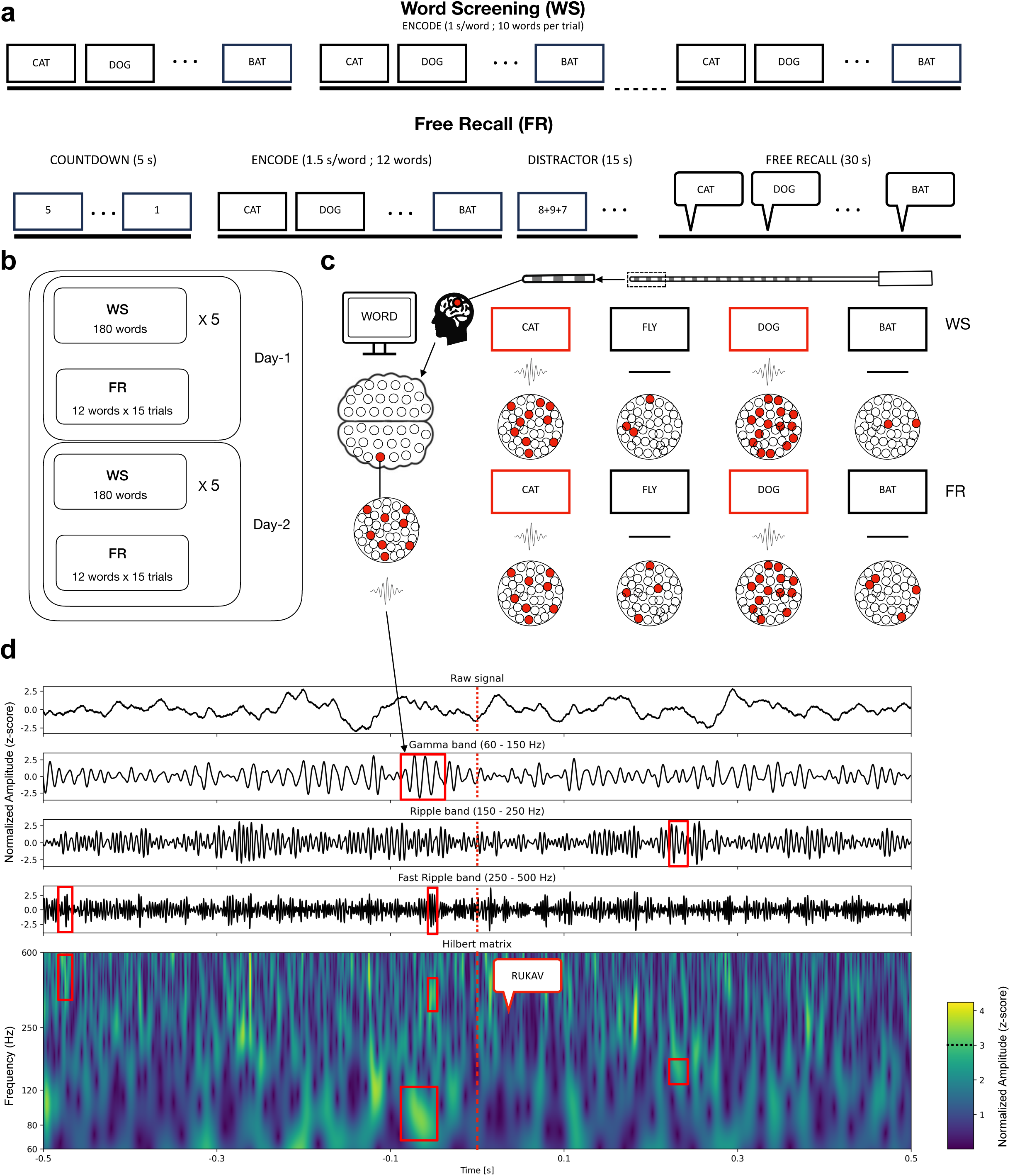
High-frequency oscillations (HFOs) induced by specific words were detected through screening and testing a pool of common nouns. (a) Schematic of two behavioral tasks used to screen and test for word-specific HFO responses. (b) Overview of the two-day study design that used the same word sets repeated across different sessions and tasks to test persistent HFO responses. (c) Diagram shows macro-contact electrode recordings sampling hypothetical local neuronal assemblies (red nodes) generating HFO bursts in response to their ‘preferred’ (red) but not the ‘non-preferred’ (black) words repeated in different tasks. (d) Example raw intracranial EEG signal (top) is decomposed into gamma, ripple, and fast ripple frequency bands to mark discrete bursts of HFOs (red rectangle) in the filtered signals and corresponding spectrogram centered around the onset of recalled word vocalization (red dashed line). Notice that each isolated burst lasts approx. four oscillation cycles with distinct peak amplitude in a particular frequency range.

**Table 1.** Summary of the patient languages, tasks and explored cortical areas. *Demographic and recording details of subjects included in the study, categorized by language (ISO 639-1 code; CS: Czech, SK: Slovak)*.

| Subject | Language<br>(ISO 639-1) | FR runs | WS runs | Macro<br>contacts | Explored<br>lobes | Age | Sex | Handedness |
| --- | --- | --- | --- | --- | --- | --- | --- | --- |
| 1 | SK | 1 | 0 | 168 | F, L | 38 | M | R |
| 2 | CS | 1 | 1 | 173 | F, L, O, P, T | 31 | F | R |
| 3 | CS | 1 | 1 | 118 | L, T | 40 | M | L |
| 4 | CS | 1 | 1 | 174 | F, L, O, P, T | 29 | M | R |
| 5 | SK | 1 | 1 | 173 | F, L, O, P, T | 21 | M | R |
| 6 | CS | 2 | 2 | 170 | F, L, P, T | 36 | M | R |
| 7 | SK | 1 | 1 | 157 | F, L, O, P, T | 31 | M | R |
| 8 | SK | 1 | 1 | 174 | F, L, T | 29 | M | R |
| 9 | CS | 2 | 2 | 172 | F, L, O, P, T | 26 | F | L |
| 10 | SK | 2 | 2 | 172 | F, L, P, T | 36 | F | R |
| 11 | CS | 0 | 1 | 137 | F, L, O, T | 30 | M | R |
| 12 | CS | 1 | 1 | 102 | F, L, O, P, T | 43 | M | R |
| 13 | CS | 1 | 2 | 134 | F, L, T | 41 | M | R |
| 14 | CS | 2 | 2 | 146 | F, L, P, T | 39 | F | R |
| 15 | SK | 1 | 1 | 159 | F, L, P, T | 37 | M | R |
| 16 | SK | 1 | 1 | 111 | L, P, T | 47 | F | R |
| 17 | CS | 1 | 1 | 137 | F, L, O, P, T | 29 | M | R |
| 18 | CS | 2 | 1 | 138 | F, L, O, P, T | 31 | F | R |
| 19 | SK | 1 | 1 | 135 | F, L, P | 32 | M | L |
| 20 | SK | 1 | 1 | 141 | F, L, O, P, T | 31 | F | R |
| 21 | SK | 1 | 1 | 162 | F, L, O, P, T | 32 | F | R |
| 22 | CS | 1 | 1 | 108 | F, P | 41 | F | R |
| 23 | CS | 1 | 1 | 84 | F, T | 27 | M | R |
| 24 | CS | 1 | 1 | 172 | F, L, O, T | 35 | F | R |
| 25 | CS | 1 | 1 | 152 | F, L, T | 33 | M | R |
| 26 | CS | 1 | 1 | 174 | F, L, O, P, T | 36 | F | R |
| 27 | CS | 1 | 1 | 128 | F, L, O, P, T | 35 | F | R |
| 28 | SK | 1 | 1 | 158 | F, L, O, P, T | 48 | F | R |
| 29 | SK | 1 | 1 | 173 | F, L, O, P, T | 25 | M | R |
| Total | CS=17,<br>SK=12 | 33 | 33 | 4302 |  | 34.1 +/- 6.4 | 16 M,<br>13 F | 26 R, 3 L |

To test this prediction, we detected individual HFO bursts within a broad high-gamma and ripple frequency range (60-200 Hz) around the times of word presentation and recall (Fig. 1d) in both tasks using methodology from our previous studies [15,18,40]. This detection method isolates distinct bursts of increased amplitude lasting at least four oscillatory cycles [41,42] and determines their peak frequency, peak amplitude, and duration in the time-frequency domain. Crucially, this prevents the detection of broadband power increases and other non-oscillatory events [40] and facilitates the separation of physiological detections from pathological activities, such as interictal epileptiform spikes and pathological HFOs [43–46], especially in channels localized in the seizure onset zone. While some spectral overlap remains unavoidable, the task-responsive HFOs identified in this study shared spectral properties consistent with physiological HFOs related to memory and cognitive functions [14,22,23].

### Word-responsive HFOs delineate the ventral visual stream and left-lateralized semantic network

Out of 3225 contacts implanted in this study across a wide range of cortical regions (Table 2) 257 (∼ 8.0%) consistently responded to word presentation with induced HFO detections (Fig. 2a). Notably, this 3,225 is a filtered subset; the full anatomical breakdown reported in Table 2 encompasses all implanted contacts, including those located in white matter. The peak latency of this induced response and the anatomical localization of the responsive contacts confirmed the expected sequence of the visual processing stream for these high-frequency activities [15,47,48]. Specifically, the induced HFOs peaked first in the sensory visual areas of the occipital cortex (100-200 ms post stimulus) and subsequently in the association and limbic areas of the temporal, parietal and frontal lobes (Fig. 2b). We observed a relatively higher density of the induced HFO detections and corresponding responsive electrodes in the posterior visual areas of the occipital and temporal cortices (Fig. 2c).

**Figure 2.**
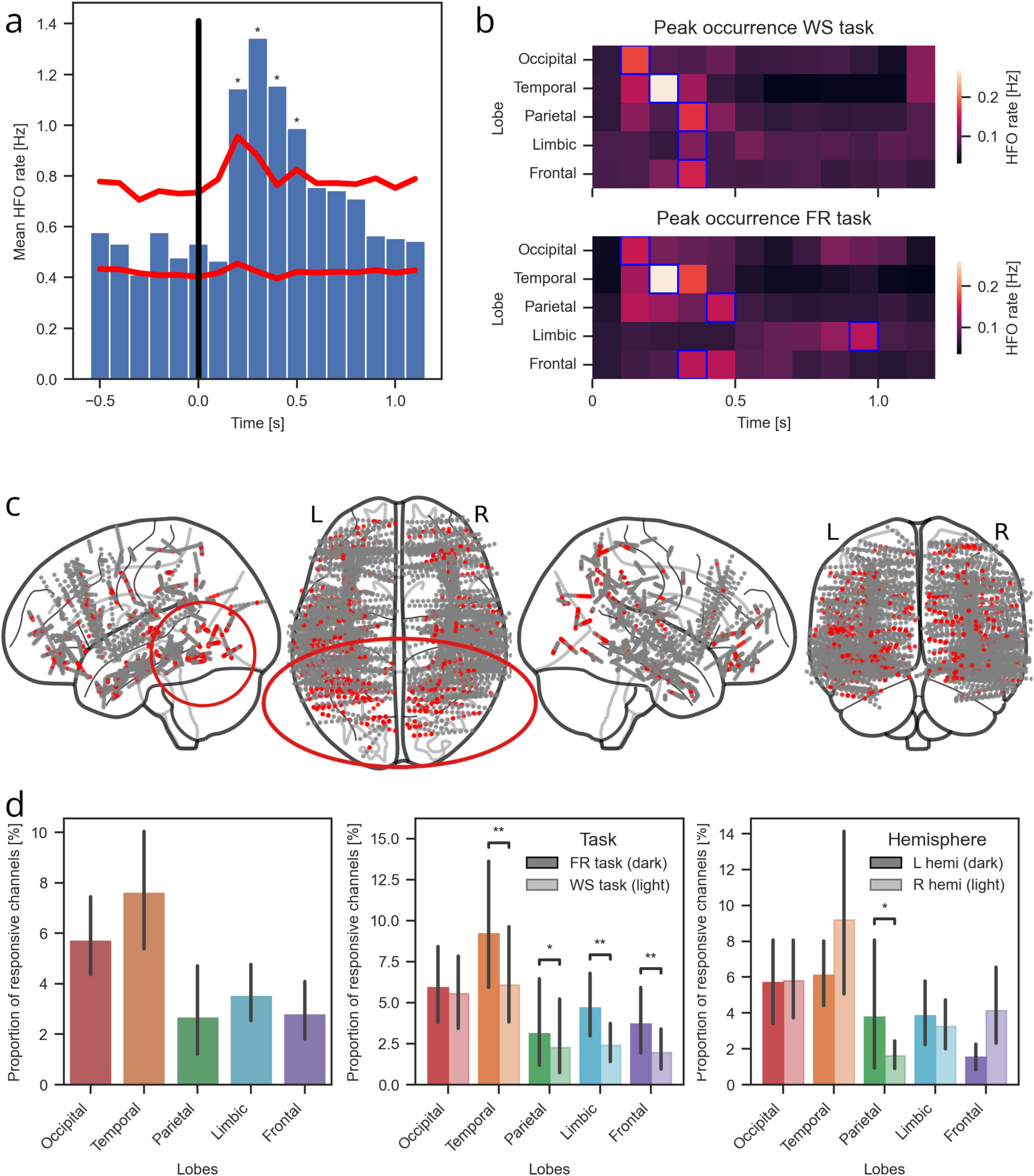
Responsive contacts with word-induced HFOs are primarily localized in the occipitotemporal processing areas. (a) Histogram of aggregated HFO detections from an example identified responsive contact shows significant responses over the baseline (red lines indicate the baseline mean and the significance threshold at mean + 3 standard deviations) following word presentation onset. (b) Heat-maps of average HFO rates from responsive electrodes show consistent peak latencies (blue square) of the visual processing stream in two tasks across the studied cortical lobes. (c) Unified brain maps illustrate localization of all identified responsive (red) and non-responsive (grey) contacts implanted. Notice dense aggregation of responsive sites (red circles) within the ventral visual processing stream. (d) Bar plots summarize the proportions of responsive contacts in the studied cortical lobes and test the effect of the screening (WS) vs memory (FR) tasks and of hemispheric laterality. Significance markers reflect a weighted mixed-effects model accounting for within-subject variance, which resolves statistical significance despite overlapping aggregate error bars (see Suppl. Fig. 1 for paired trajectories)

**Table 2.** Total counts of electrode contacts localized in distinct anatomical structures. Electrode placement varied across cortical regions based on clinical priorities, leading to non-uniform coverage across patients. The table summarizes the number of electrodes implanted per structure from all patients, reflecting variability in cortical sampling due to patient-specific clinical needs.

| Occipital lobe |  | Temporal lobe |  | Parietal lobe |  | Limbic structures |  | Frontal lobe |  |
| --- | --- | --- | --- | --- | --- | --- | --- | --- | --- |
| Structure | N contacts | Structure | N contacts | Structure | N contacts | Structure | N contacts | Structure | N contacts |
| Calcarine fissure | 21 | Fusiform gyrus | 103 | Angular gyrus | 74 | Amygdala | 71 | Gyrus rectus | 68 |
| Cuneus | 23 | Heschl's gyrus | 27 | Inferior parietal gyrus | 99 | Cingulate gyrus | 79 | Inferior frontal gyrus | 190 |
| Fusiform gyrus | 9 | Inferior temporal gyrus | 203 | Paracentral lobule | 4 | Hippocampus | 348 | Orbitofrontal cortex | 138 |
| Inferior occipital gyrus | 13 | Insula | 323 | Postcentral gyrus | 166 | Parahippocampalgyrus | 95 | Paracentral lobule | 9 |
| Lingual gyrus | 57 | Middle temporal gyrus | 686 | Superior parietal gyrus | 70 |  |  | Precentral gyrus | 83 |
| Middle occipital gyrus | 30 | Superior temporal gyrus | 240 | Supramarginal gyrus | 126 |  |  | Precuneus | 28 |
| Precuneus | 63 | Temporal Pole | 96 |  |  |  |  | Rolandic operculum | 61 |
| Superior occipital gyrus | 23 |  |  |  |  |  |  | Superior frontal gyrus | 56 |

We assessed differences in the regional distribution of responsive channels using a binomial generalized linear model (GLM), revealing a significant effect of anatomical lobe on the proportion of responsive channels relative to the total number of contacts. Subject identity was included as a covariate to account for inter-individual variability in overall channel response rates. The model revealed a significant effect of anatomical lobe on the proportion of responsive channels (Wald χ²(4) = 238.28, p = 2.172 × 10⁻⁵⁰). A significant effect of hemisphere (Wald χ²(1) = 5.92, p = 0.015) and task (Wald χ²(1) = 34.64, p = 3.960 × 10⁻⁹) were also observed. Additionally, a significant interaction between factors was detected (likelihood ratio test, p = 7.976 × 10⁻³), indicating that the effect of lobe varied across conditions. Overall, the temporal lobe exhibited the highest probability of containing responsive channels relative to other lobes (Fig. 2d). We found significant task-related effects in the temporal (z = −3.35, p = 0.002), parietal (z = −2.28, p = 0.028), limbic (z = −3.70, p = 0.001), and frontal lobes (z = −3.17, p = 0.003), whereas the occipital lobe did not show a significant task-related difference (z = −0.92, p = 0.360) (Fig. 2d). Additionally, we observed a significant lobe × hemisphere interaction (p = 0.008), indicating that hemispheric differences varied across lobes. Post hoc contrasts revealed significant hemispheric asymmetry only in the parietal lobe (z = 2.79, p = 0.026), whereas the occipital (z = 1.39, p = 0.204), temporal (z = −1.00, p = 0.317), limbic (z = 1.64, p = 0.167), and frontal lobes (z = 2.02, p = 0.108) did not show significant lateralization (Fig. 2d). Because our model isolates consistent within-subject effects across a highly variable dataset, statistical significance is captured even where aggregate group-level confidence intervals visually overlap (Fig. 2d; see Suppl. fig. 1 for detailed within-subject effects). These results confirm a prediction that the visual areas of the occipital cortex would be equally engaged in sensory processing of the presented stimuli irrespective of cognitive demands of a given task. In all other association areas, the induced HFOs reflected a higher engagement when the same words were presented to be remembered in the memory task than just attended during word viewing in the screening task.

More detailed analysis of the anatomical localization of the implanted electrode contacts in particular structures within the studied lobes revealed a more stratified and diverse distribution of the responsive contacts (Suppl. fig. 2). Notably, the highest proportions of these contacts were localized in the ventral visual stream, including the calcarine, fusiform, and lingual gyri, which are involved in processing simple and complex features of visual word forms. When evaluating task-related engagement, significant effects of the memory versus the screening task (Suppl. Fig. 3) were identified in the hippocampus (z = 1.05, p = 0.017), parahippocampal gyrus (z = −0.31, p = 0.038), and orbitofrontal cortex (z = 0.52, p = 0.017)—structures known to be central to the mesial temporal lobe and limbic memory systems. Furthermore, after controlling for multiple comparisons, we found significantly more responsive channels in the inferior frontal gyrus (z = 4.17, p = 4.48 × 10⁻⁴) and the middle temporal gyrus (z = 4.41, p = 3.03 × 10⁻⁴) of the left hemisphere (Suppl. Fig. 4). These are known to be key structures for language processing within semantic cortical networks [48]. Overall, these findings confirm that the HFO responses were induced selectively more in the areas associated with verbal and declarative memory processing in the human brain.

### HFO word selectivity increases along a posterior-to-anterior cortical gradient

Having determined the responsive electrode contacts in our studied tasks, we estimated how many words from the pool these contacts respond to. Given that macro-contacts sample from a wide population of neuronal assemblies, we expected any single contact to respond to multiple words (see Fig. 1c). We identified ‘preferred’ words by comparing the high-frequency oscillation (HFO) detections induced by five repetitions of a given word in the WS task against those of all other words (Fig. 3a). For each channel, we evaluated whether individual words elicited a significant response. Across all sessions, this produced 45,714 word–channel observations (180 words × number of channels). Of these, we classified 8,177 (17.9%) as preferred words.

**Figure 3.**
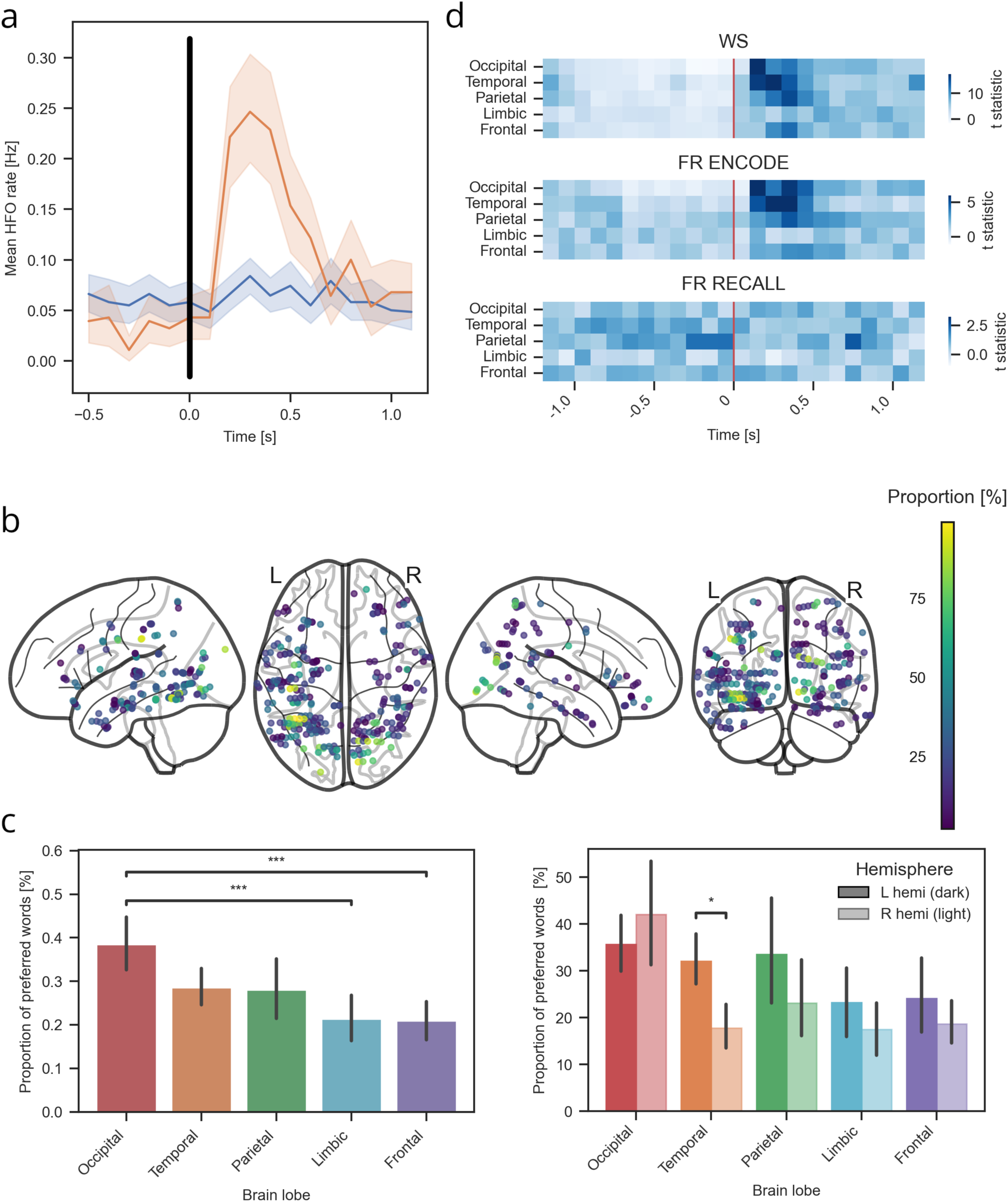
Specificity of HFO responses to preferred words is gradually increasing along the cortical processing hierarchy. (a) Example of preferred word HFO detections from an identified responsive contact. (b) Unified brain maps of word-specificity of all identified responsive contacts reveals a posterior-to-anterior trend of gradually decreasing proportions (warm colors) in higher-order processing areas. (c) Bar plots confirm significantly lower proportions of preferred words and significant laterality effects in the anterior compared to the posterior occipital area.(d) Heatmaps display average t-statistics from all preferred word contact responses aligned to the onset of word presentation (WS and FR encode) and recall vocalization (FR recall). Notice relatively higher t-statistic values right after word presentation and before recall onset despite no external stimulation in the latter. * – p<0.05, ** – p < 0.01, *** – p < 0.001.

This proportion of preferred words differed across the cortical lobes, revealing a consistent anatomical trend (Fig. 3b). Unified brain maps of word-selectivity revealed a posterior-to-anterior trend of gradually decreasing proportions of preferred words in higher-order processing areas. We observed the highest proportion in the posterior sensory regions of the occipital lobe (39.3%), followed by gradually lower proportions in the successively more anterior association regions of the temporal (28.7%), parietal (22.4%), frontal (21.0%), and limbic cortices (10.3%). The same trend of increasing word selectivity was also observed at the level of individual cortical structures (Suppl. Fig. 5). Contacts located in early visual sensory regions responded, on average, to approximately 40% of the presented words. In contrast, contacts situated in higher-order associative areas exhibited markedly greater selectivity, typically responding to only ∼10–20% of the words. This pattern suggests a gradual increase in response selectivity along the sensory to higher-order association cortical areas. Latency of the preferred-word responses in each lobe followed by this visual processing stream hierarchy (Fig. 3b).

We found a significant effect of cortical lobe on the proportion of the preferred words (Wald χ2(4) = 19.20, p = 7.182 × 10⁻⁴). Pairwise post-hoc comparisons revealed a higher proportion of preferred words in the occipital compared to the limbic (z = 3.880, p = 1.046 × 10⁻⁴) and frontal (z = 3.757, p = 1.720 × 10⁻⁴) lobes, and a similar trend compared to the temporal (z = 1.889, p = 0.0589) and parietal (z = 1.678, p = 0.0933) lobes (Fig. 3c). Post-hoc contrasts revealed a significant hemispheric difference only in the temporal lobe (p = 0.013), whereas the occipital (p = 0.539), parietal (p = 0.505), limbic (p = 0.578), and frontal lobes (p = 0.594) did not show significant lateralization (Fig. 3c).

### Preferred-word responses persist across tasks, memory phases, and consecutive days

Finally, we examined the temporal dynamics of these responses across different brain structures and time bins using event-aligned heatmaps, where the t-statistic quantified the difference in activity between preferred and non-preferred words (Fig. 3d). We observed relatively higher t-statistic values immediately following word presentation during the encoding phases (WS and FR). Notably, significant activity was also observed just before recall vocalization in the FR task, despite the absence of external stimulation during that period.

Finally, we addressed the key question of this study – are these HFO responses to particular preferred words preserved across (1) encoding and recall of the same task, (2) across the two tasks, and (3) across repeated sessions on two subsequent days? Having determined the preferred words for each contact in the WS task, we tested whether the same words will induce significantly more HFOs during the FR encoding and recall phases within that Day 1 session and in the WS and FR phases on the next Day 2 session (Fig. 4 top left). In the original WS task on Day 1, we found high selectivity of HFO responses to preferred words in posterior temporal and occipital lobe regions, including the fusiform gyrus (FFG), inferior temporal gyrus (ITG), middle temporal gyrus (MTG), superior temporal gyrus (STG) and lingual gyrus (LING), as well as in the parietal, limbic and frontal lobe structures at latencies corresponding to visual information processing of the presented words (Fig. 4). These spatiotemporal patterns of selective HFO responses were closely replicated during word presentation in FR task. What is more, the preferred-word responses persisted also to the FR recall phase, in which they were observed even before the onset of word verbalization with no external sensory stimulation (Fig. 4 top right). The same anatomical structures still showed significant responses on the next day, especially in a subset of posterior cortical lobe structures (Fig. 4 bottom), although the spatiotemporal patterns were less conserved across the two days than within the same original Day 1 session mainly due to a limited number of subjects with two sessions of each task completed (Table 1).

**Figure 4.**
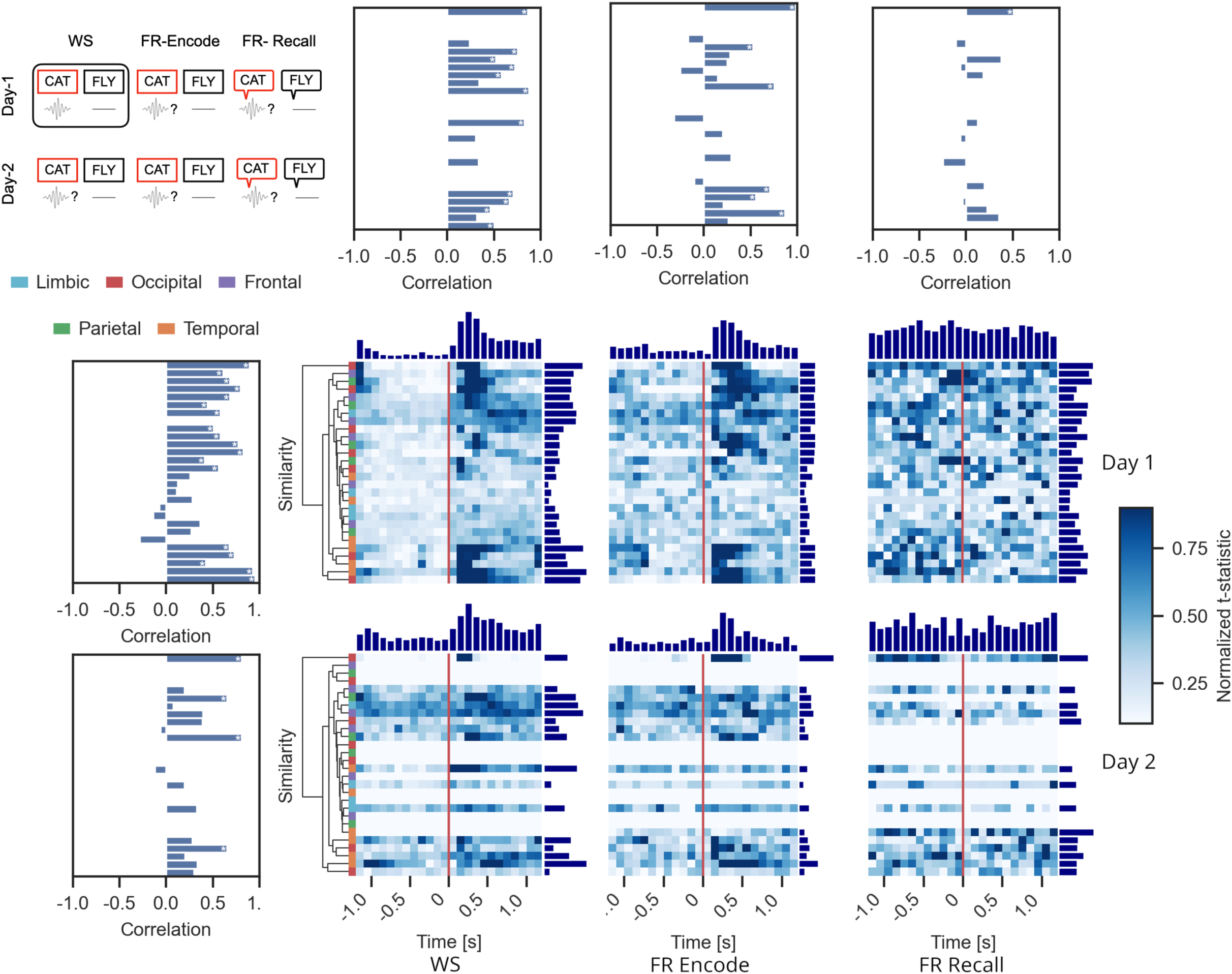
Word-specific HFO responses are preserved across tasks, memory phases, and days. (Top left) Diagram of the experimental design testing whether preferred words identified during the first screening (WS – black rectangle) would selectively induce HFOs to the same words presented or freely recalled in other tasks on the same and on the subsequent day. (Center and bottom right) Heat-maps quantify differences in the average preferred word responses across all responsive contacts localized in particular cortical structures and in time bins (column plots summarize the aggregate values for the anatomical area and the time bin) aligned to the onset of word presentation or recall vocalization. Notice persistent preferred-word responses among hierarchically clustered brain structures (color bands indicate cortical lobe membership) even before free recall vocalization with no external cuing or stimulation. (Top and left) Bar-plots summarize correlation of temporal response profiles between WS and FR task phases on a given day (left) and within the same task phase across consecutive days (top) in any one cortical structure. Notice multiple significant correlations (white asterisks) for most structures across the tasks on Day 1 that are persisting but gradually fading across the three phases (horizontally) and across subsequent days (vertically) from the original WS screening of the identified preferred words. * – p<0.05

We found strong positive correlations (Pearson’s correlation, p < 0.001) between encoding phases of the two tasks (Fig. 4 left) in multiple visual and ventral temporal structures, including lingual gyrus (r = 0.95), inferior temporal gyrus (r = 0.93), fusiform gyrus (r = 0.89), cuneus (r = 0.83), and superior occipital gyrus (r = 0.80), among other significant correlations in parietal and prefrontal areas. Limbic and orbitofrontal regions showed a weak cross-task similarity. A similar pattern was observed across recording days (Fig. 4 top). During WS encoding, response profiles were highly consistent between days in ventral temporal and parietal structures, including the fusiform gyrus (r = 0.86, p = 5.819 × 10⁻⁸), supramarginal gyrus (r = 0.87, p = 2.487 × 10⁻⁸), insula (r = 0.83, p = 6.143 × 10⁻⁷), middle temporal gyrus (r = 0.71, p = 1.128 × 10⁻⁴), and middle occipital gyrus (r = 0.67, p = 3.720 × 10⁻⁴). Comparable stability was observed for FR encoding, particularly in the fusiform gyrus (r = 0.98, p = 2.808 × 10⁻¹⁸), inferior temporal gyrus (r = 0.87, p = 4.457 × 10⁻⁸), supramarginal gyrus (r = 0.75, p = 2.291 × 10⁻⁵), and middle temporal gyrus (r = 0.71, p = 1.156 × 10⁻⁴). In contrast, correlations across days during FR recall were generally weaker and reached significance only in the fusiform gyrus (r = 0.51, p = 0.012). Overall, the early sensory areas exhibited the strongest cross-task and cross-day similarities, whereas higher-order association areas showed greater variability in their response profiles.

### Hierarchical spatiotemporal clustering reveals global networks of semantic processing

The word-specific HFO responses were found at distinct latencies from word presentation, depending on their anatomical location. Hence, we clustered them according to temporal similarity of the responses between the studied structures (see Fig. 4). The resultant spatiotemporal clusters revealed a sequence of temporally coordinated networks from multiple cortical lobes. We found a clear temporal order of the word-specific responses, spanning the entire word presentation period from early visual to higher-order semantic information processing (Fig. 5a). Each cluster of the studied structures peaked at a distinct time (Fig. 5b), which generally corresponded to anatomical location in the ventral visual processing stream; clusters with earlier (short latency) peak of the word-specific HFO responses comprised mainly occipitotemporal cortical areas, whereas the ones with late peaks engaged more anterior temporal and prefrontal areas (Fig. 5c). However, there was a considerable proportion of contacts in the posterior regions of the fusiform gyrus that participated together with the anterior regions in the late latency clusters, suggesting higher-level language processing also in these visual areas [49]. The general posterior-to-anterior continuous sequence of word-specific HFO responses is reminiscent of the hierarchical order of induced high frequency spectral activities [15,48] and their subsequent memory effects [47] that were previously reported in similar studies.

**Figure 5.**
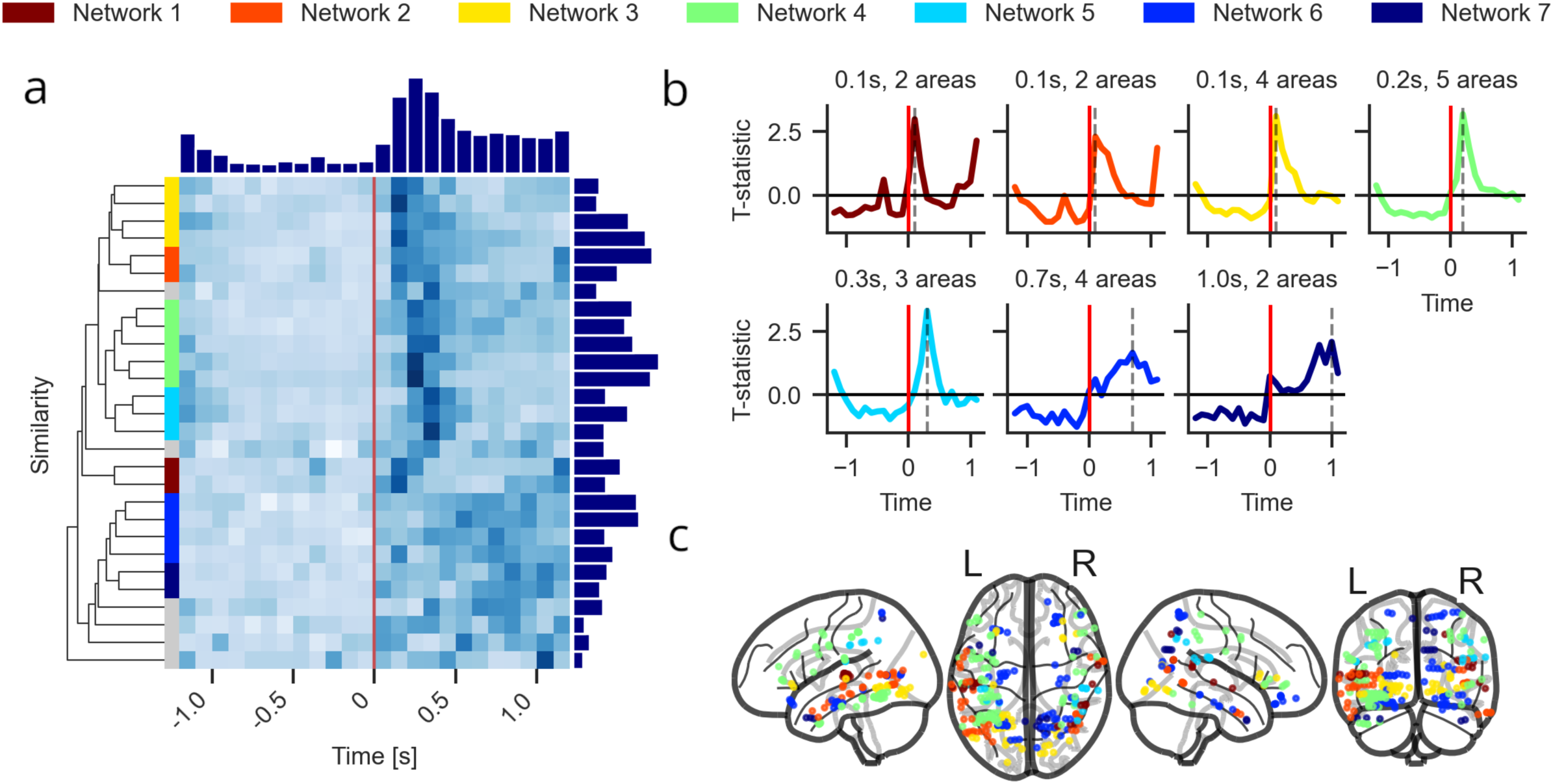
Hierarchical spatiotemporal clustering of word-specific HFO responses reveals global networks of visual and semantic information processing. (a) Heat-map of normalized temporal profiles of the preferred-word HFO responses (plotted as in Fig. 4) shows global networks between the studied cortical structures (color-coded) ordered by the peak response latency. Notice the descending amplitude profile of the preferred-word responses (top and right column plots) from the onset of word presentation (horizontal) and from the top network clusters (vertical). (b) Line plots (smoothed) summarize the average temporal profiles of the identified network clusters (color-coded and ordered as in ‘a’). Notice the continuous sequence of the peak latencies starting early right after word presentation through to late toward the end of the presentation period. (c) Anatomical distribution of the responsive contacts on the unified brain maps visualizes widespread distribution of the identified color-coded network clusters. Notice that the early and late peak clusters are not limited to the posterior or the anterior cortical areas, respectively, but reveal mixed sensory visual and higher-order semantic area localizations.

## Discussion

In this work, we tested a hypothesis that HFOs detected on clinically used intracranial macro-electrode contacts reflect the processing of information about specific word concepts. Our results showed that on average approx. every tenth contact responded to word presentations with an increased rate of HFO detections. HFOs were induced in response to a subset of words from the tested pool, depending on the anatomical localization of the contact along the ventral visual processing stream. The lowest word selectivity (i.e. the largest subset of preferred words) was found in the posterior sensory visual and the highest (i.e. the smallest subset of preferred words) in the anterior associational cortical areas. In all the areas, the identified contacts responded selectively to their preferred words across different memory phases, tasks and sessions of testing. These persistent word-specific HFO responses were induced at distinct times of presentation on the screen and before recall vocalization with no sensory stimulation, revealing a sequence of global cortical networks involved in visual and semantic processing of word concepts. We propose that the HFO responses to specific preferred words on any one electrode contact reflect the underlying electrophysiological activities of multiple neuronal assemblies, corresponding to particular concept representations.

To begin with, our study was based on a hypothesis that HFOs detected on macro-contacts can be used to sample coordinated activities of neuronal assemblies underlying word concepts. Previously, we have shown that gamma and ripple frequency HFOs detected on macro-contacts are associated with memory encoding and recall [15,40]. Compared to micro-contact recordings of single neuron activities that can encode specific concepts [2,3,5,8,50], macro-contact recordings of HFOs from large populations of neuronal assemblies were first shown to be expectedly less selective and respond to more general features of the encoded stimuli [31].

However, subsequent studies linked high gamma/ripple frequency HFOs detected on the macro-contacts with encoding and recall of specific memory traces [17,18,25,28,32,34]. Combined micro– and macro-contact recordings showed that single neuron firing underlying ripple frequency HFOs can predict successful recall of specific word concepts [28]. In particular, the relative sequence of single-unit spiking between neuronal assemblies that are engaged in HFO generation contains information about specific encoded and recalled stimuli [27,51,52]. Whether micro-scale information from neuronal spiking can be detected and decoded from emergent macro-scale HFOs remains underexplored.

Hence, we proposed HFOs as large-scale electrophysiological biomarkers of neuronal assemblies to track their activities on the micro-, meso– and macro-scales of cortical organization [14,22]. They are detected across a large scale of neural activities, ranging from the canonical bands of the EEG spectrum (1-60 Hz) up to the high frequency ranges (60-600 Hz) using macro– and micro-electrode contacts [21,24]. We hypothesized that an individual HFO burst sampled from a macro-contact reflects an emergent discharge of a particular neuronal assembly that processes information about a specific word concept. Multiple assemblies recorded within one macro-contact sampling field would give rise to several HFOs induced by presentation of their ‘preferred’ words (see Fig. 1). If true, then one should detect more HFOs in response to the preferred words that are processed by neuronal assemblies within the macro-contact sampling field. The preferred word responses would also be expected to persist across different tasks and time, assuming that the assemblies were stable in their spatiotemporal dynamics. Effectively, these are the criteria for dynamic engram activities of neuronal assemblies [35,36,53] that persist from formation to retrieval of sensory and higher order representations for particular stimuli.

We tested these hypothetical assembly activities at the level of induced HFO responses across sensory and higher order processing areas. The highest density of stimulus-specific responses was observed in the visual and semantic areas of the ventral visual stream. The relative proportion of word-responsive contacts was the highest in the posterior occipitotemporal areas of the visual cortex and was progressively smaller in more anterior semantic areas of the frontal and limbic areas. The proportion of preferred words that these contacts responded to follow the same posterior-to-anterior gradient of increasing selectivity, i.e. HFO responses to gradually smaller proportions of words (see Fig. 2-3). These results are in agreement with the previous similar studies of induced spectral power in the high frequency ranges [15,47,48]; higher trial-averaged spectral power observed in the visual areas can be explained on a trial-by-trial level by relatively higher proportion of ‘preferred’ stimuli that these more sensory areas responded to as compared to more selective association areas in the anterior temporal, parietal, limbic and prefrontal cortex. Given that the more anterior areas were localized in the semantic system of the brain [54], they were likely responding to gradually more specific word concepts.

We expected this gradual refinement or tuning of stimulus-specific responses along the ventral visual processing stream. Neurons at the top of this stream in the limbic perirhinal-hippocampal areas are known to process increasingly complex and unique stimuli [55]. Concept cells provide an extreme example of tuning and selectivity for multimodal representations of a single object [7,50]. This highly specific neuronal firing has recently been associated with ripple frequency HFOs in the limbic structures [29,30]. The limbic areas in our study showed the highest selectivity, i.e., the lowest proportion of the preferred stimuli, of around 10% of all words from the pool. Compared to the micro-scale single neuron recordings, this is still a relatively high selectivity given the macro-scale recordings of HFOs that are generated by multiple neuronal assemblies from the sampled population. Apart from the highly specific concept cells, there are other single unit cell types in the limbic structures that respond to more general categories and features shared by multiple stimuli [8,56,57]. We found single units in the same patient dataset as used in this study, which respond to approx. 5-10% of the presented words (data not shown). In general, our results show that macro-scale HFO response selectivity ranges from 10-40% depending on the anatomical localization in the processing stream. Selectivities below 10% would be expected for single unit activities or possibly also micro-contact HFO recordings, which remain to be tested. Low selectivity above 40% of the presented stimuli were observed in the same dataset for global coincident HFO bursting across multiple cortical areas [18]. By and large, our findings suggest stimulus selectivity depends both on the anatomical localization and on the scale of the electrode recordings. The selectivity of our macro-scale HFO responses places between more specific single-unit micro– and less selective macro-scale recordings of global coincident HFO bursting [18].

In contrast to the classic paradigms investigating encoding and recall selectivity in one task, we used the same pool of words administered in a battery of different tasks repeated on subsequent days [38,39]. The tasks probed attention/immediate recall (WS), free recall (FR) and cued recall (PAL). There were visual and auditory versions of the FR and PAL tasks administered in four languages with automated transcription [58] but only WS and FR tasks had enough data to be included in this study (see Table 1). Having both visual and spoken expressions of the same words in multiple languages applied in the context of various cognitive tasks repeated over subsequent days, provides ideal conditions for testing stability and generalizability of the specific stimulus representations. The preferred word responses determined in the WS task visual presentations were highly conserved in the equivalent FR task presentations, but also before and during verbal recall of the same words with no visual presentation (see Fig. 4). It means that recall of abstract concept representations of the preferred words in the mind even before they were vocalized was associated with selectively more HFOs than recall of the other non-preferred words. This concept-specific HFO activity started as early as 1200 ms before the onset of vocalization (no visual stimulation) and was observed on the contacts in both sensory visual and semantic areas, suggesting abstract, multimodal word representations that emerged in the subject’s imagination. How conscious and aware subjects were of these representations is another question that our paradigm did not address. One recent study linked hippocampal sharp-wave ripples with the emergence of spontaneous, self-generated thoughts [59]. Other studies proposed coincident cortical HFOs in binding coherent perceptual and mnemonic representations [16–18,33]. Our results confirm that the word-specific HFO responses persisted across different tasks, modalities, and time. It remains to be determined whether the emergent representations can be decoded using HFOs to identify specific concepts triggered by external sensory stimulation or internal self-generated free recall.

If so, HFOs could provide a feasible substrate for tracking electrophysiological engram activities [22]. An activity that is induced by external stimulation and by internal cuing of a memory trace would fulfill the requirements for an engram activity. In our study, viewing the presented words in the WS task or encoding and recalling them in the FR task would require retrieving specific engrams for particular words. Therefore, similar neuronal assemblies should be reactivated in each of these various conditions. Our results showed persistent spatiotemporal patterns across the studied anatomical structures and the time of word presentation or free recall (see Fig. 4-5), which is reminiscent of pattern reinstatement between encoding and retrieval of specific stimuli across multiple cortical areas [60–62]. Similarity in these spatiotemporal patterns revealed clusters of anatomical structures forming global networks across different cortical lobes. For example, one such cluster with an early latency peak of preferred-word responses comprised contacts from the occipital and temporal lobes, whereas another with a later latency peak comprised prefrontal and temporal cortex contacts (see Fig. 5). Interestingly, the latter included both higher-order semantic processing areas in the prefrontal cortex as well as more sensory areas of the fusiform gyrus. Such global networks would explain higher-level language processing even in the occipitotemporal cortex at early stages of the ventral visual processing stream [49]. Engrams for word concepts would necessarily engage widespread networks from multiple lobes that support their multimodal sensory aspects and semantic meanings. Global coincident HFOs have recently been proposed as a feasible mechanism to support the functioning of such widespread networks and the emergent perceptual and/or memory representations [16,18]. Similar mechanisms of global integration of information processing were proposed for functions involving subjective consciousness [63], suggesting a more universal role of the HFO activities that extend beyond engrams and memory functions. Although it remains speculative at this point, HFOs offer a testable substrate for memory and other higher-order cognitive functions.

Using macro-scale HFO detections to study cognitive functions in this patient population has several limitations. First of all, interictal epileptiform activities are detected with the same electrode contacts in similar frequency ranges. Even though pathological and physiological HFOs have overlapped spectral properties [43,45], frequency-at-peak and amplitude can be used to dissociate them. Fast ripple frequency ranges beyond 250 Hz are more likely to be pathological based on characteristic single-unit activity patterns [64]. Still, separating pathological discharges or non-oscillatory artifacts from physiological HFO detections remains a major challenge [14,22,23]. Even classic physiological hippocampal sharp-wave ripple complexes require development of more advanced, feature-based detection and classification methods to properly separate them from pathological events [65]. In our study, the spectral properties of each detection were carefully determined (see Fig. 1) and used as inclusion criteria for physiological HFOs [44]. To further exclude a potential influence of epilepsy-related activities, we designed the methodology to detect HFOs induced by cognitive events associated with memory processing like word presentation or recall. Pathological discharges are less likely to be consistently induced by such cognitive events and hence outnumbered by physiologically induced HFOs. Cognitive stimulation may actually suppress and effectively lower the probability of pathological HFOs [43,66]. Overall, while the presence of pathological activity cannot be fully excluded from these intracranial macro-contact recordings, their potential impact can be minimized.

Due to these precautions, we took a conservative methodological approach to carefully screen and select only those contacts that consistently respond to word presentations. This prevented us from including potentially pathological responses to single words but limited us from detecting physiological responses that were selective for few words. Therefore, the proportions of responsive contacts and the preferred word selectivity could have been higher, but we deliberately intended our analysis to be minimally affected by potential pathological activities. Given that the criteria for physiological and pathological HFOs or for oscillatory and non-oscillatory activities have been a subject of a long-standing debate, we chose to be on the conservative side at the expense of losing some true physiological detections. This was part of the reason also for limited data availability from particular anatomical structures (see Table 2) on the Day 2 sessions (see Fig. 4). Our study was limited to only two days of recordings during tasks, which provided only a partial view of processing specific stimuli. Continuous chronic recordings over multiple days of online task performance and offline quiet wakefulness and sleep will be inevitable for testing the stability and selectivity of the macro-scale HFO responses for particular concepts.

To conclude, in this work we presented proof-of-concept for macro-scale HFO responses to specific common nouns that are independent from sensory stimulation, cognitive tasks, or the time of testing. Testing beyond the timeframe of two subsequent days remains to be performed with other verbal and non-verbal modalities, e.g., spoken words or their images. Tracking such multimodal representations across time and anatomical areas will require large-scale recordings from multiple high-density electrode arrays [67–70] to determine the source of HFO activities. Micro-electrode recordings could resolve the source activities and spatial volume of individual HFO bursts on the level of single cortical columns [22]. How widely spread and dynamic are the underlying neuronal assemblies and neural networks? How do they change in the process of learning, short– and long-term memory, consolidation, or development? What is the difference between online and offline processing of their preferred stimuli? Can they be targeted and modulated to treat memory and cognitive deficits at the fundamental level of specific engrams? Emerging neurotechnology for concurrent high-density recording and stimulation of HFO activities are ideally suited to address these pending questions.

## Methods

### Participants and Electrode Contact Localization

Twenty-nine patients (16 males; mean age 34.1 ± 6.4 years) with pharmacoresistant epilepsy undergoing intracranial stereo-EEG (sEEG) monitoring for epilepsy surgery at St. Anne’s University Hospital in Brno were recruited for this study (Table 1). Participants were native speakers of Czech (ISO 639-1: CS; n = 17) or Slovak (ISO 639-1: SK; n = 12); Table 1). All participants provided written informed consent prior to participation. The study protocol was approved by the local Institutional Review Board and conducted in accordance with the Declaration of Helsinki. The trajectory and placement of depth electrode probes were determined exclusively by clinical requirements for seizure localization, resulting in variable cortical coverage across participants (Suppl. Table 1).

Penetrating depth electrodes (AdTech Inc., PMT Inc., DIXI Inc.) were implanted. Reference signals were obtained via additional subgaleal electrodes. Post-implantation, pre-operative high-resolution CT scans were co-registered with post-operative MRI scans. Electrode contact locations were normalized to MNI space and anatomically labeled using the Automated Anatomical Labeling atlas version 3 (AAL3)[71]. For analysis, contacts were grouped into five primary regions of interest: frontal, parietal, temporal, occipital, and limbic regions. The limbic category included the hippocampus, amygdala, parahippocampal cortex, and posterior cingulate cortex.

### Experimental design

Participants completed a battery of cognitive tasks targeting eye movements, semantic processing, and verbal memory. In the present study, we analyzed two verbal memory paradigms: Word Screening (WS) and Free Recall (FR). In the Free Recall (FR) task, participants were presented with a list of 12 words displayed sequentially on a computer screen. Each word was drawn from a pool of common nouns shared with other verbal memory tasks in the experimental battery. Participants were instructed to memorize the words for subsequent recall. Following the presentation of the word list, a brief distractor phase was administered to prevent rehearsal. The distractor task consisted of simple algebraic equations that participants were required to solve. After completion of the distractor phase, participants were asked to recall as many words as possible from the previously presented list, in any order, by vocalizing them freely.

The Word Screening (WS) task was administered at the end of each experimental session. During the task, participants were instructed to attend to words presented sequentially on a computer screen. The stimuli consisted of 180 unique words drawn from the same word pool used in the FR task. To ensure sustained attention, participants were intermittently prompted to repeat the most recently presented word aloud. This prompt was indicated by the appearance of three question marks on the screen. One WS trial consisted of the presentation of a full set of 180 words. Participants completed five trials in total, such that each word was presented five times over the course of the task. The WS task was designed to elicit word-specific neural responses independent of explicit memory demands, allowing identification of cortical regions selectively responsive to individual lexical items or concepts.

The FR and WS tasks were repeated on a subsequent day using identical word lists presented in the same fixed order. This design enabled the examination of neural correlates of encoding, maintenance, and retrieval, as well as cross-day reinstatement of electrophysiological activity patterns. Due to clinical constraints, not all patients completed the second session (see Table 1).

### Electrophysiological Recordings

Intracranial EEG (iEEG) signals were acquired using the BrainScope BioSDA09 system (M&I Ltd.), with up to 192 channels recorded per subject at a sampling rate of 5,000 Hz. Signals were recorded from standard depth electrodes with a 2.3 mm exposed contact surface and 5–10 mm inter-contact spacing. During acquisition, a subgaleal electrode served as a reference. For analysis, signals were re-referenced using a bipolar montage between adjacent contacts to improve spatial specificity and reduce common-mode artifacts. All signals were visually inspected.

Because all participants were patients with drug-resistant epilepsy, recordings may contain pathological activity such as interictal spikes or high-frequency oscillations (HFOs). However, several factors mitigate their influence on the present analyses: (i) the event-related design attenuates non-time-locked pathological activity through averaging, (ii) epileptiform discharges are often suppressed during cognitive engagement, and (iii) analyses focused on frequencies below 200 Hz, avoiding pathological fast ripples. To ensure transparency and data portability, all electrophysiological recordings were formatted in MEF 3.6 and structured in accordance with BIDS protocols. All data were de-identified to protect participant privacy in adherence to established ethical guidelines. Access to the dataset is provided through the EBRAINS repository [39]

### Detection of individual HFO bursts

To ensure high spatial specificity and minimize volume conduction artifacts, signals from neighboring contacts on each depth electrode were bipolar-referenced. HFOs were then identified using an automated detection pipeline adapted from our earlier work [15,18,40]. First, the referenced signals were decomposed into 38 logarithmically spaced frequency bins between 60 and 500 Hz using zero-phase finite impulse response filters (200 ms transition bands, −6 dB cutoff). The amplitude envelope for each frequency band was extracted utilizing the Hilbert transform and subsequently normalized via a sliding z-score transformation, calculated using moving 10-second baseline windows.

To delineate the start and end boundaries of putative HFOs, candidate events were initially marked when the z-scored envelope exceeded a liberal threshold of 2 standard deviations (SD) for a minimum of three consecutive oscillatory cycles. Concurrent detections spanning multiple frequency bands were merged into unified events. For each consolidated event, the peak amplitude and its corresponding peak frequency were assigned based on the single data point exhibiting the highest z-score.

To differentiate true physiological HFOs from pathological epileptiform activity or artificial spectral power increases (e.g., resulting from filtering sharp signal transitions), a stricter set of secondary criteria was applied. Final HFO detections were restricted to the 80–200 Hz range. These candidate bursts were only retained if they achieved a peak amplitude greater than 3 SD and persisted for more than 3 complete oscillatory cycles at the peak frequency (Figure 1d). Additionally, a final cycle-count verification confirmed that at least one full oscillation occurred strictly above the threshold. This detection protocol was executed within a defined time window of −1.5 s to +1.5 s relative to either stimulus onset (word presentation) or the beginning of vocalization during the free recall task.

### Identification of Responsive Channels

Responsive channels were identified by analyzing the temporal distribution of high-frequency oscillation (HFO) detections across trials. The 2.4-second window was divided into non-overlapping 100 ms time bins. For each channel, HFO counts were summed across all word presentations. Channel selection was based exclusively on the encoding phase of the tasks.

To determine consistent task-related responsiveness, a cross-channel statistical thresholding approach was applied independently to each time bin. For every 100 ms bin, the mean and standard deviation of trial-summed HFO counts were calculated across all channels. A channel was classified as “responsive” if its HFO count in any bin exceeded the global cross-channel mean plus three standard deviations (mean + 3 SD). This conservative threshold ensured the selection of channels demonstrating robust time-locked responses across multiple stimulus presentations while maximizing the signal-to-noise ratio for subsequent analyses. Responsive channel determination was performed separately for the Word Screening (WS) and Free Recall (FR) tasks.

### Anatomical Distribution of Responsive Channels

To examine whether responsive channels were preferentially distributed across cortical regions, we quantified the proportion of responsive channels relative to the total number of implanted contacts within each anatomical structure. Differences across lobes were evaluated using a generalized linear model (GLM) with binomial error distribution and logit link function. The dependent variable represented the number of responsive channels relative to the total number of contacts (responsive contacts / total contacts). Lobe, hemisphere, and task were included as fixed effects, alongside their interaction terms, to assess condition-specific anatomical effects. Unequal numbers of implanted contacts across structures were incorporated through binomial weights. A similar approach was used to evaluate the distribution of responsive channels across cortical structures. Structure, hemisphere, and task were included as fixed effects, and interaction terms between structure and hemisphere, and between structure and task, were included to assess structure-specific effects. Subject identity was included as a covariate to account for inter-individual variability, and unequal numbers of implanted contacts were incorporated through binomial weights.

### Identification of Preferred Words

Following identification of responsive channels, we examined whether these channels exhibited word-specific HFO responses during the Word Screening (WS) task. For each responsive channel, HFO counts were analyzed in 100 ms non-overlapping bins across the same −1.2 to +1.2 second event window. A pre-stimulus baseline window from −500 ms to 0 ms relative to stimulus onset was used to estimate baseline HFO activity for each channel.

In each responsive channel a word was classified as “preferred” if the HFO count in any post-stimulus bin (0 to +1.2 s) exceeded the baseline mean plus three standard deviations (baseline mean + 3 SD). Words not meeting this criterion were categorized as non-preferred. Preferred words were defined exclusively using data from the first day of the WS task, establishing a reference dictionary used for independent analyses of the Free Recall task.

To assess regional differences in selectivity, the proportion of preferred words relative to the full WS word pool was computed for each channel and aggregated across anatomical regions.

Differences in preferred-word selectivity across anatomical regions were evaluated using generalized estimating equations (GEE) with a binomial distribution and logit link function. The dependent variable represented the proportion of preferred words relative to the total number of words presented for each channel. Anatomical region and hemisphere were included as predictors, and subject identity was treated as the clustering variable to account for correlations among channels recorded within the same participant. An exchangeable working correlation structure was assumed. Hemisphere effects within individual lobes and structures were assessed using linear contrasts of model coefficients.

### Cross-Task Reactivation Analysis

To investigate the reinstatement of word-specific responses during memory encoding and retrieval, HFO detections in the Free Recall task were analyzed using identical filtering parameters and temporal binning (100 ms bins).FR encoding trials were time-locked to the onset of word presentation on the screen, whereas FR recall trials were aligned to the onset of the participant’s vocalization. Both phases were analyzed within a −1.2 to +1.2 second window relative to the corresponding event.

Reactivation strength was quantified by comparing HFO responses elicited by preferred and non-preferred words. For each channel and anatomical structure, HFO counts were contrasted within each time bin, and reactivation strength was expressed as the t-statistic comparing responses to preferred versus non-preferred words (Figure 4). To assess the stability of structure-specific response profiles across experimental conditions, Pearson correlation coefficients were computed between temporal response vectors across tasks (WS vs FR) and across recording days (day 1 vs day 2). Because recall responses were temporally shifted relative to stimulus onset, mean t-statistics across time bins were computed separately for encoding and recall phases prior to correlation analysis.

### Structure-Level Analysis of Temporal HFO Response Profiles

To characterize temporal response dynamics across cortical structures, t-statistics were normalized by z-scoring across time bins within each structure. Hierarchical clustering was applied to these normalized temporal profiles to identify groups of structures exhibiting similar response dynamics (Figure 5a). Cluster determination was performed using agglomerative linkage on the normalized time-resolved t-statistic vectors. The optimal number of clusters was determined using the elbow method by examining the within-cluster sum of squares as a function of cluster number and selecting the point of diminishing returns, discarding clusters that included only one structure (N=1). Cluster membership was visualized by plotting mean temporal profiles for each cluster and by mapping cluster assignments onto the spatial distribution of implanted electrode contacts (Figure 5b. 5c).

### Statistical Analysis

All statistical analyses were performed in Python using the StatsModels library. Differences in anatomical distributions of responsive channels were evaluated using generalized linear models (GLMs) with a binomial error distribution. In these models, Lobe, Task (Word Screening vs. Free Recall), Hemisphere, and their interactions were included as fixed effects, while subject identity was included as a covariate. Regional differences in word selectivity, as well as task-related activation effects at the level of individual fine-scale anatomical structures, were analyzed using generalized estimating equations (GEEs) to account for correlated measures, with subject identity treated as the clustering variable. The statistical significance of main effects and interactions in the GLM and GEE models was evaluated using Wald χ2 tests. Where main effects were significant, pairwise post-hoc comparisons between lobes were conducted using linear contrasts. Similarity of structure-specific response profiles across conditions was assessed using Pearson correlation coefficients. Multiple comparisons across structures, time bins, and post-hoc contrasts were controlled using the Benjamini–Hochberg false discovery rate procedure (FDR correction). All tests were two-sided with a significance threshold of α = 0.05.

## Data and Code Availability

The Human brain local field potential recordings during a battery of multilingual cognitive and eye-tracking tasks (v1) data used in this study are available in the EBRAINS database at https://doi.org/10.25493/4FZH-ZCG. All analyses were conducted on data from this openly accessible resource. The raw LFP data are protected and are not available in the public repository due to data privacy laws regarding human participants. The processed data used in this study are available at EBRAINS. The code for the analysis is available in the following GitLab repository: https://gitlab.com/brainandmindlab/memory_encoding.

## Acknowledgment

This research was fully supported by the National Science Centre, Poland, grant Opus LAP number: 2020/39/I/NZ4/02070, by the Czech Science Foundation (GAČR), project No. 21-44843L. Preliminary results for this study were supported by the Ulam NAWA grant awarded to Jan Cimbalnik. The study would not be possible without a dedicated effort of the patients and their families. The authors would like to thank Betul Celik, Karolina Kacprzycka and Dimitris Panagiotis for their student project results, which helped in the design and interpretation of this study.

## Author contributions

Conceptualization: MTK, JC, SP

Data Curation: LJ, SP, MK, PD, RR, MTK, JC.

Formal Analysis: JC, SP, MTK.

Funding Acquisition: JC, MTK.

Investigation: MTK, SP, JC.

Methodology: JC

Project Administration: MTK, JC.

Resources: MTK, JC.

Supervision: MTK

Validation: SP, MTK, JC.

Visualization: SP, JC.

Writing – Original Draft Preparation: JC, SP, MTK.

Writing – Review & Editing: JC, SP, LFSR, LJ, MK, PD, RR, AC, MB, MTK.

## Tables

**Supplemental Table 1.** Summary of clinical profiles and seizure-related data. Clinical information for all participants, including dataset identifiers, seizure onset zone (SOZ) channels and corresponding anatomical structures, MRI findings, and postoperative seizure outcome when available (Engel classification).

| ID | SOZ channels | SOZ structures | MRI Findings | Engel Score |
| --- | --- | --- | --- | --- |
| 1 | B'1-2, C'1-2 | Hippocampus | Normal | IA |
| 2 | - | - | Normal | - |
| 3 | B1-4, C1-3, B'1-5, C'1-3 | Hippocampus, Parahippocampal gyrus | Normal | - |
| 4 | G'1-2, R'1-6, S'1-3 | Cingulate gyrus, Precuneus | Normal | IVB |
| 5 | G'1-4, H'7-11 | Middle temporal gyrus | Normal | IV |
| 6 | Y'2-10, I'1-4 | Insula | Venous angioma, cavernoma parieto-occipital left | IV |
| 7 | I'1-6 | Paracentral lobule, Postcentral gyrus | Polymicrogyria postcentral sulcus left | IIIA |
| 8 | - | Middle frontal gyrus, Superior frontal gyrus | Normal | - |
| 9 | B1-3, C1-3, B'1-3, C'1-5 | Hippocampus, Parahippocampal gyrus | Normal | - |
| 10 | Y'1-4, I'1-5 | Insula | Normal | - |
| 11 | O5-7 | Fronto-occipital cortex | Posttraumatic changes temporal right |  |
| 12 | B'1-4, C'1-4 | Hippocampus | Posttraumatic changes temporo-occipital left | - |
| 13 | Y1-4, X1-3 | Insula | Normal | - |
| 14 | B1-2 | Parahippocampal gyrus | Postsurgical changes right anterior mesial temporal resection (AMTR) | - |
| 15 | - | - | Normal | - |
| 16 | B1-4, C1-3 | Right hippocampus | Hippocampal sclerosis | - |
| 17 | B1-4, C1-3 | Temporal operculum | Normal | - |
| 18 | T3-6, B1-2 | Left hippocampus and amygdala | Normal | - |
| 19 | A'1-4, B'1-2 | Left hippocampus | Hippocampal sclerosis | - |
| 20 | B'1-2, C'1-2 | Fronto-parietal left |  | - |
| 21 | network epilepsy | Temporal pole |  | - |
| 22 | D1-6 | Right dorsolateral prefrontal cortex | Postoperative changes frontal right | - |
| 23 | W3-4 | Left dorsolateral prefrontal cortex | Postoperative changes frontal left | - |
| 24 | Q5-10 | Left hippocampus, lateral temporal cortex | Normal | - |
| 25 | P'1-5 | - | Postoperative changes temporal right | - |
| 26 | B'1-4, C'1-4, P'4-9, T'1-5, C'8-12, A'5-11, B'10-14 | - | - | - |
| 27 | 06-10, C1 | Parietal cortex | - | - |
| 28 | - | Right hippocampus | - | - |
| 29 | - | - | - | - |

**Supplementary Figure 1.**
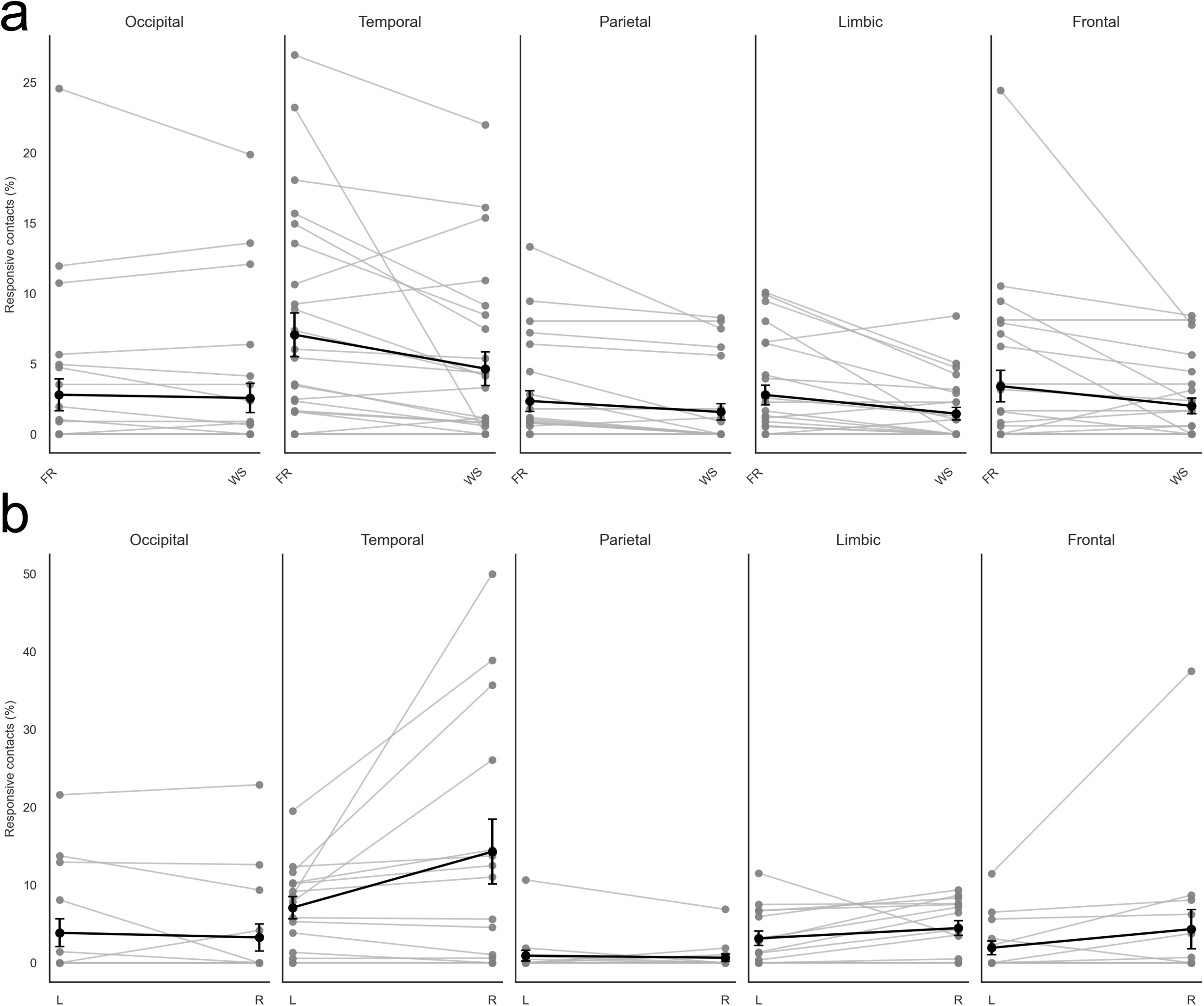
Paired individual subject trajectories of responsive contacts. Plots display the percentage of responsive contacts across cortical lobes comparing (a) task conditions (Free Recall [FR] vs. Word Specific [WS]) and (b) hemispheres (Left [L] vs. Right [R]). Individual subject data (gray lines) and group means (black lines ± SEM) highlight the consistent within-subject differences driving the statistical significance of the generalized linear model, despite substantial inter-subject baseline variability.

**Supplemental Figure 2.**
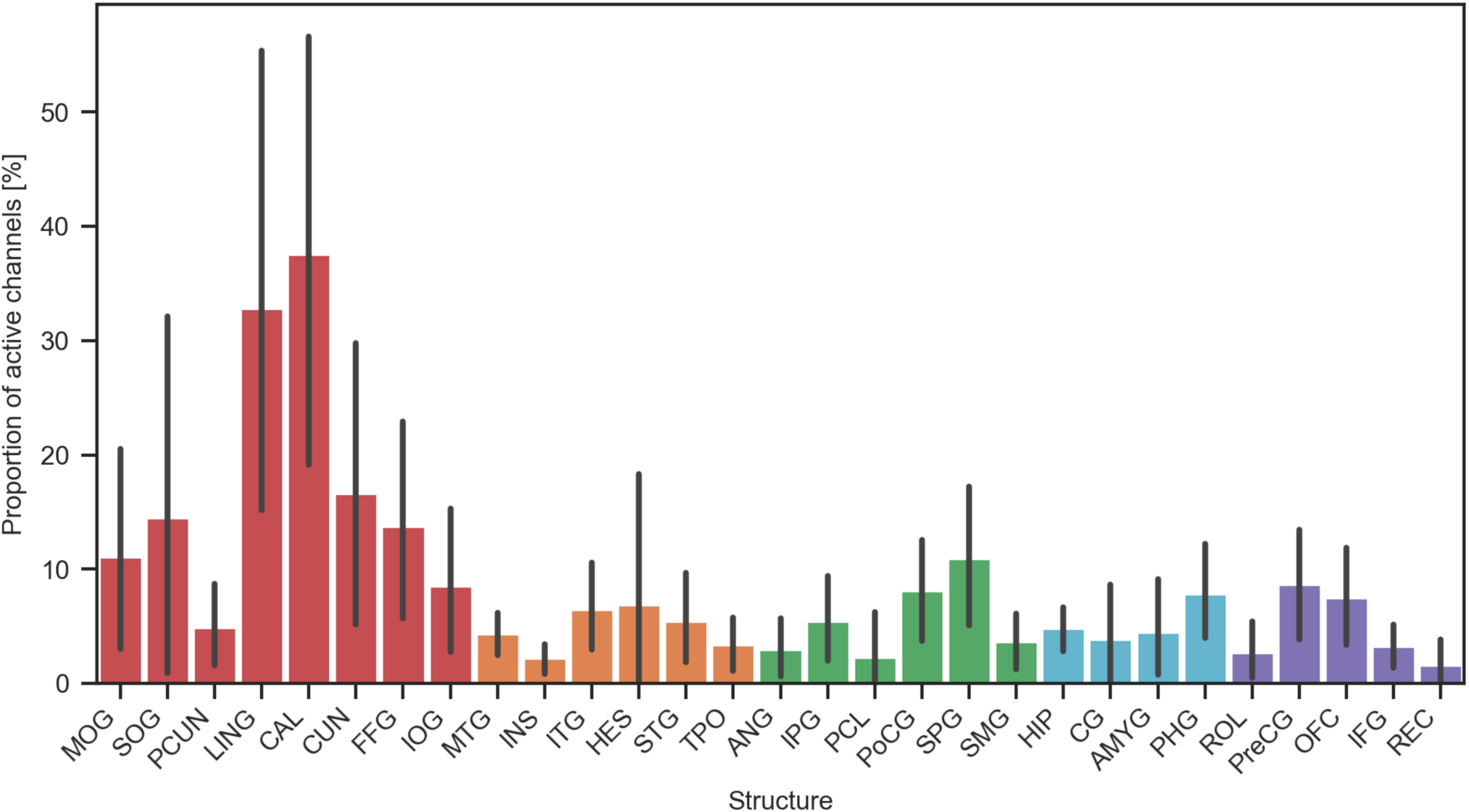
Distribution of responsive channels across cortical structures. Bar plot summarizes the proportions of responsive contacts in the studied cortical structures and tests the effect of the screening (WS) vs memory (FR) tasks and of hemispheric laterality. * – p<0.05, ** – p < 0.01, *** – p < 0.001.

**Supplemental Figure 3.**
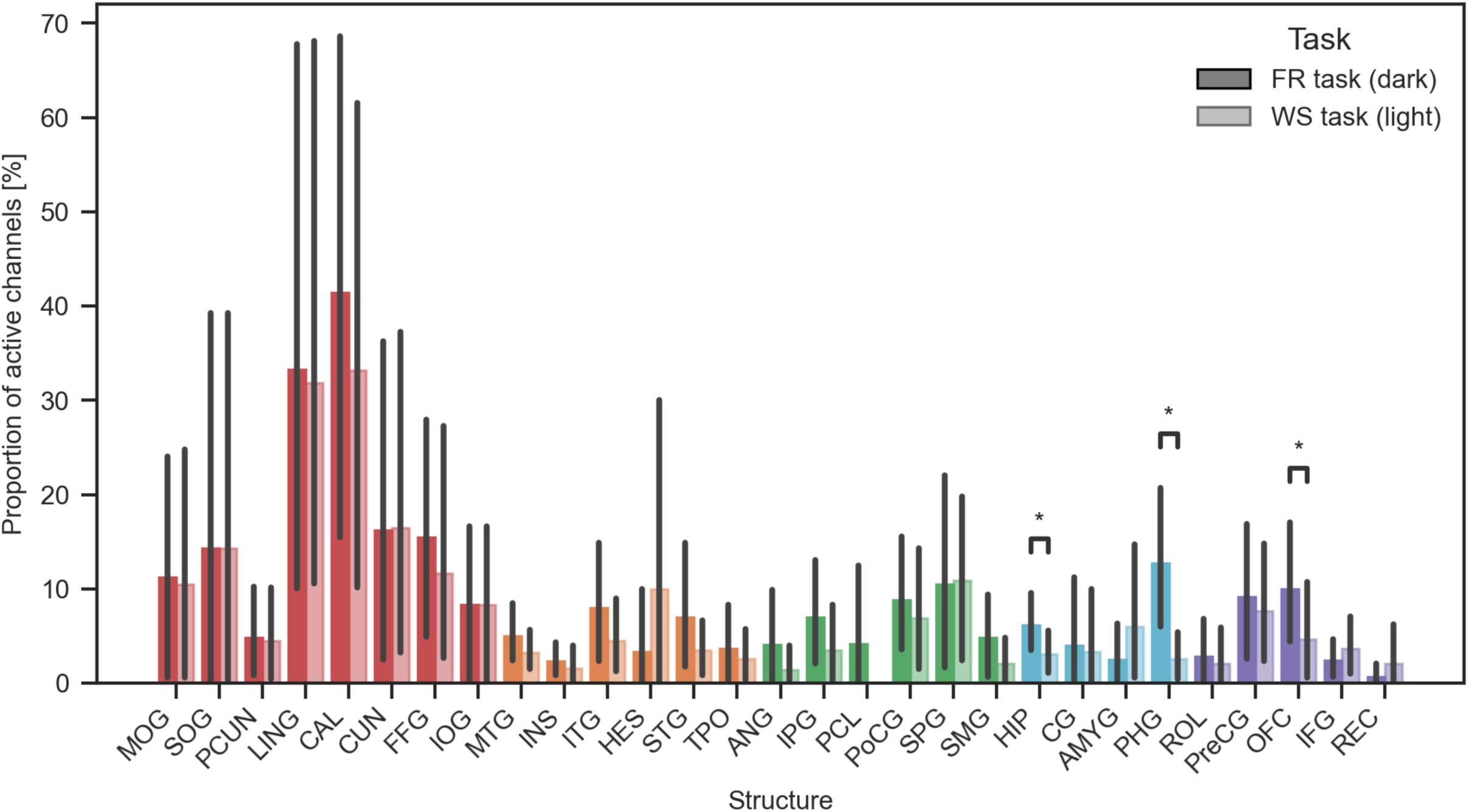
Distribution of responsive channels across cortical structures with regard to task. Bar plot summarizes the proportions of responsive contacts in the studied cortical structures and test the effect of the screening (WS) vs memory (FR) tasks. * – p<0.05, ** – p < 0.01, *** – p < 0.001.

**Supplemental Figure 4.**
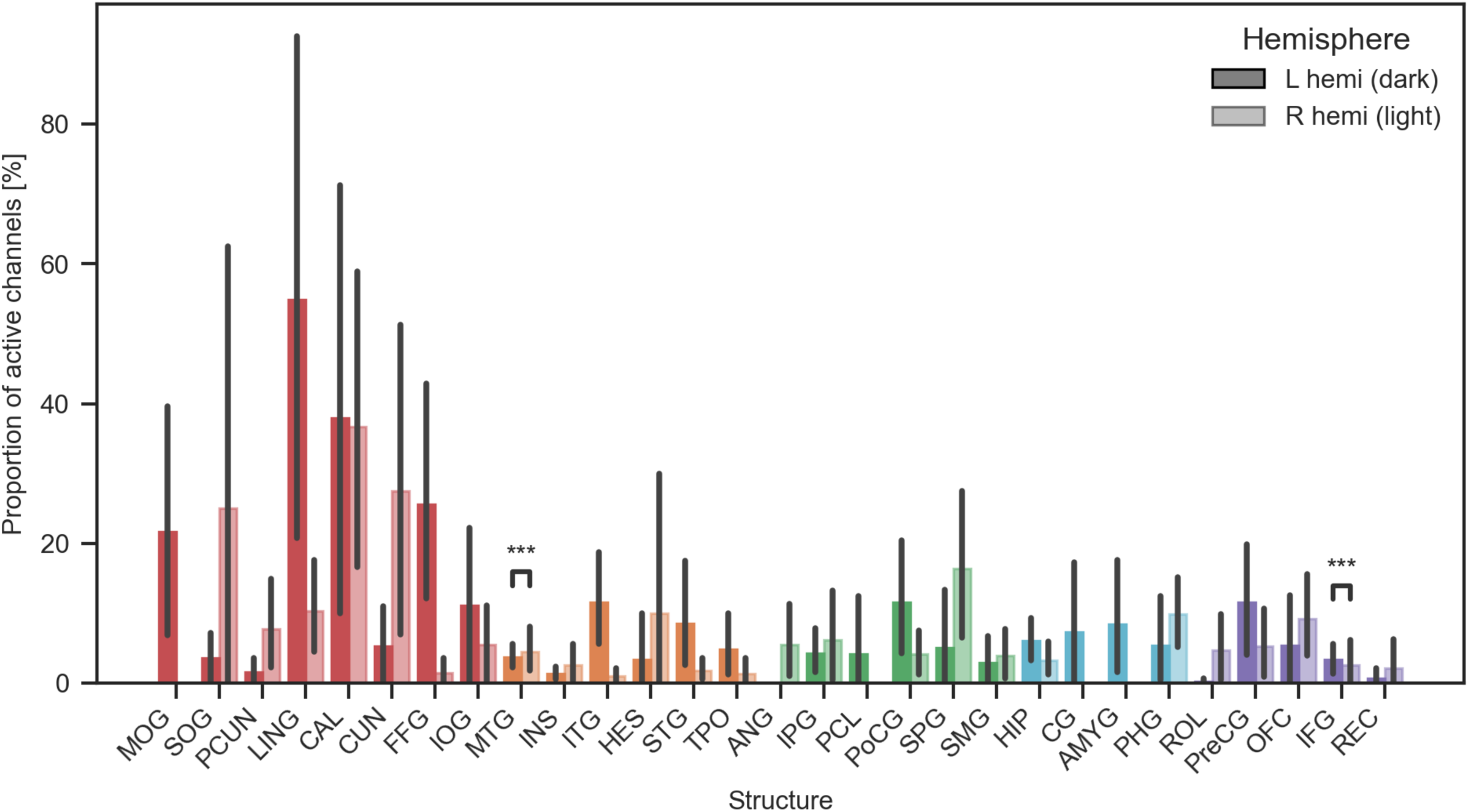
Distribution of responsive channels across cortical structures with regard to hemispheres. Bar plot summarizes the proportions of responsive contacts in the studied cortical structures and tests the effect of hemispheric laterality. * – p<0.05, ** – p < 0.01.

**Supplemental Figure 5.**
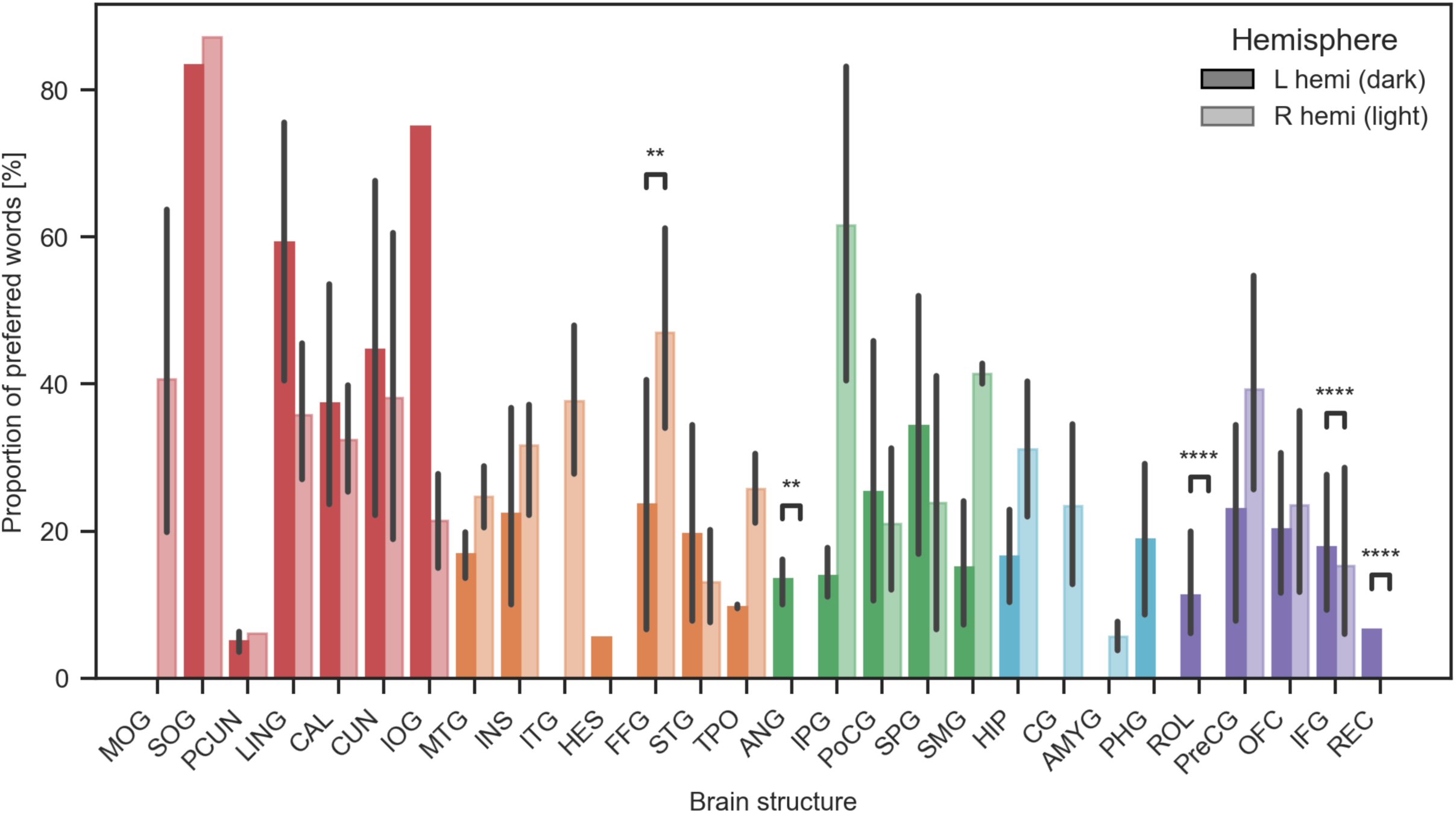
Distribution of preferred words across cortical structures with regard to hemispheres. Bar plot summarizes the proportions of preferred words in the studied cortical structures and tests the effect of hemispheric laterality. * – p<0.05, ** – p < 0.01, *** – p < 0.001.

